# RBP4-BACH1 Interaction Modulates Transcriptional Regulation of Insulin Signaling Pathway Genes

**DOI:** 10.64898/2026.08.24.746665

**Authors:** Li Wang, Qi Ma, Yao Chen, Chunmei Wu, Baoping Guo, Mayire Nuermaimaiti, Yinxia Su, Binbin Fang, Lijuan He, Aliya Rehati

**Author notes:** **Corresponding Authors:** Lijuan He, Fax: + 86/991/4361202, Aliya Rehati, Fax: + 86/991/4361202. These authors contributed equally to this work (Li Wang, Qi Ma, Yao Chen).

## Abstract

Retinol binding protein 4 (RBP4) exhibits diurnal oscillatory pattern and is elevated under conditions of circadian disruption and in type 2 diabetes mellitus, yet the molecular link between RBP4 and impaired glucose metabolism remains elusive. Here, we overexpressed RBP4 in human hepatoma Huh7 cells and performed integrated RNA sequencing (RNA-seq), Co-immunoprecipitation (Co-IP) coupled with mass spectrometry (MS), and Cleavage Under Targets and Tagmentation (CUT&Tag). We identified BACH1 as a direct RBP4□interacting transcription factor that predominantly binds the TGACTCA motif in promoter regions of genes involved in carbon metabolism pathways. Integrative analysis of RNA seq and CUT&Tag data uncovered 63 direct target genes co regulated by RBP4 and BACH1, including known circadian and metabolic regulators SLC7A11, PFKFB3, CTCF, NR1D2 and WEE1 as well as novel candidates SF1 and PIN1. These target genes are significantly enriched in insulin receptor signaling and carbohydrate metabolic pathways. Mechanistically, the RBP4–BACH1 axis reprograms glucose metabolism, linking circadian rhythm disturbances to dysregulated glucose homeostasis. Collectively, our findings establish a functional role for RBP4 in connecting circadian disruption to diabetes and highlight RBP4 as a potential therapeutic target.

## Introduction

Diabetes has emerged as a major global public health challenge. The global prevalence of diabetes among adults aged 20–79 years reached 589 million in 2024 and is projected to rise to 853 million by 2050(Federation, 2025). The incidence of diabetes is strongly correlated with obesity, metabolic disorders, and lifestyle changes. Recent studies have demonstrated that disrupted circadian rhythms constitute a critical trigger for the onset of T2DM. Irregular sleep patterns(Chaput *et al*, 2024), night-time light exposure(Windred *et al*, 2024), and shift work(Shan *et al*, 2018) significantly increase the risk of diabetes, indicating that the hepatic circadian clock is a principal driver of rhythmic gene expression in glucose and fatty acid metabolic pathways. Nevertheless, the mechanisms through which circadian rhythms regulate hepatic metabolic homeostasis via molecular networks, thereby influencing the pathological progression of T2DM, remain incompletely understood.

RBP4 is not only involved in retinol metabolism, but also closely related to insulin resistance, T2DM(Fan & Hu, 2024), obesity(Sandoval-Bórquez *et al*, 2024), cardiovascular and cerebrovascular risk(Bustamante *et al*, 2021) and other metabolic diseases. Our preliminary findings revealed that RBP4 expression levels are elevated in both shift workers and individuals with diabetes(Li *et al*, 2023), suggesting that RBP4 might serve as a molecular bridge between circadian rhythm and T2DM. As a adipokine, RBP4 interacts with the STRA6 receptor, which is the main known mechanism by which it regulates insulin sensitivity(Gliniak *et al*, 2017). However, it is unclear whether RBP4 directly regulates gene expression in liver cells such as interacting with transcription factors in addition to this classical signaling pathway. And if there is other non classical pathways also need to be explored. Notably, circulating RBP4 predominantly originates from the liver, a central metabolic organ. The circadian regulatory functions of the liver are pivotal in the development and progression of T2DM. Liver-specific knockout of circadian clock genes has been shown to induce insulin resistance and abnormal gluconeogenesis(David *et al*, 2015). However, the key effector molecules mediating these processes remain unidentified. We therefore hypothesize that RBP4 may serve as a critical molecular bridge linking circadian rhythms to T2DM.

Furthermore, although the role of the transcription factor BACH1 in iron metabolism(Fuminori, 2024) and oxidative stress(Xiangxiang *et al*, 2025) has been partially characterized, its interaction with RBP4 and its specific functions in the liver circadian rhythm–metabolism axis remain largely unexplored. Despite these clues, there is still a lack of data on the role and regulatory mechanisms of RBP4 at the transcriptional level.

The present study aims to investigate the mechanisms through which RBP4 interacts with transcription factors to regulate metabolism. Using Huh7 cells as a model system, Co-IP coupled with MS was performed to identify transcription factors that bind to RBP4, which had been previously screened by RNA-seq. Subsequently, CUT&Tag analysis was performed to identify BACH1-bound DNA targets. Integration of RNA-seq data from RBP4-overexpressing cells revealed that the co-regulated target genes of RBP4 and BACH1 were significantly enriched in the insulin receptor signaling pathway and carbohydrate metabolism processes. This study demonstrates, for the first time, that RBP4 participation in hepatic circadian rhythm and metabolic homeostasis through a BACH1-mediated transcriptional regulatory network. These findings provide a novel perspective on the pathogenesis of T2DM and establish a theoretical foundation for developing therapeutic strategies targeting the circadian rhythm–metabolism axis.

## Results

### 2.1 Identification of BACH1 as a Key Transcription Factor Directly Interacting with RBP4

Based on our previous findings(Li *et al*., 2023) suggesting that RBP4 may serve as a molecular link between the circadian rhythm and T2DM, RNA-seq analysis (GSA-Human: HRA016389) was performed following RBP4 overexpression and NC in Huh7 cells to identify differentially regulated genes. These results led us to hypothesize that RBP4 might interact with specific TFs. To test this hypothesis, Co-IP followed by MS (GSA-Human: HRA016411) was conducted in Huh7 cells to capture RBP4-associated proteins. The input group (Input) served as a baseline control representing background levels, whereas the IgG-immunoprecipitated (IgG) group functioned as a negative control. Notably western blot analysis of Co-IP confirmed that, in two independent replicate experiments, the target protein RBP4 with approximately 23 kDa was detected in the RBP4-immunoprecipitated group (IP) (Figure 1A).

**Fig 1.**
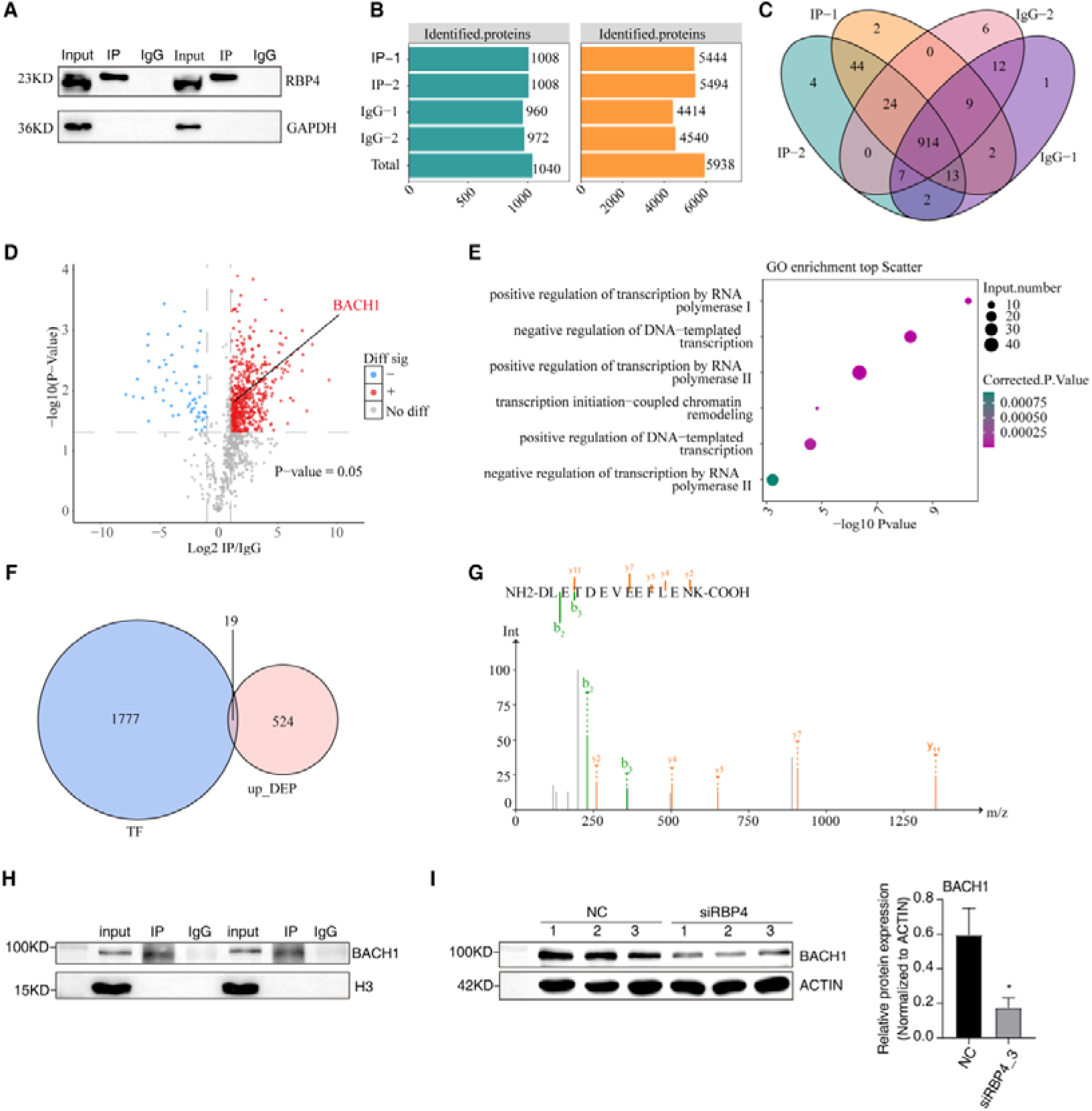
RBP4 interacts with BACH1 in Huh7 cells. (A) RBP4 protein detection by western blot in combined CO-IP. (B) The number of proteins identified (left) and the number of peptides identified (right). The abscissa represents the statistical number, and the ordinate represents the number of samples. (C) Venn diagram showing the protein intersection between different samples. (D) The volcano plot shows RBP4 interacting proteins, upregulated in red and downregulated in blue. (E) GO function enrichment analysis of the genes corresponding to RBP4 interacting protein. (F) Mass spectrometry identified BACH1 peptides. (G) The secondary spectrum of BACH1. (H) Co-IP/WB to detect RBP4-BACH1 interaction. (I) WB analysis of BACH1 protein expression after RBP4 knockdown.

The IP and IgG samples from two biological replicates each were subsequently subjected to mass spectrometry sequencing. In total, 5,938 peptides and 1,040 proteins were identified across the four samples (Figure 1B). Overlap analysis revealed that 995 proteins were consistently detected in both IP replicates (44 + 24 + 914 + 13). Compared with the IgG control samples, 50 proteins were specifically enriched in the IP group (2 + 44 + 4) (Figure 1C). Differential quantitative analysis was then conducted on the identified proteins, and the volcano plot highlighted in red the proteins interacting with RBP4. Among the 543 proteins identified as RBP4 interactors, BACH1 was specifically annotated (Figure 1D).

To elucidate the functional and biological relevance of these RBP4-interacting proteins, GO enrichment analysis was performed (Figure 1E). The most significantly enriched pathways included positive regulation of transcription by RNA polymerase I, negative regulation of DNA-templated transcription, positive regulation of transcription by RNA polymerase II, transcription initiation-coupled chromatin remodeling, and both positive and negative regulation of DNA-templated transcription. Given our focus on TFs interacting with RBP4, the list of identified genes was intersected with known TF gene sets, resulting in 19 candidate TFs potentially interacting with RBP4 (Figure 1F). Based on the roles of RBP4 in circadian rhythm-related metabolism and BACH1 in hepatic insulin signaling, BACH1, which was prominently highlighted in the proteomic profile, was selected for further validation. The secondary mass spectrometry spectrum of the characteristic peptide segment demonstrates the specificity and accuracy of the captured BACH1 (Figure 1G). Western blotting of MS validated the interaction between RBP4 and BACH1. BACH1 expression was detected in Input and IP groups while no target protein was detected in IgG groups, indicating an interaction between BACH1 and RBP4 (Figure 1H). Furthermore, RBP4 knockdown in Huh7 cells led to a reduction in BACH1 expression, as evidenced by Western blot analysis (Figure 1I). Collectively, these findings demonstrate a binding interaction between RBP4 and BACH1 in Huh7 cells.

### 2.2 BACH1 Regulated Metabolic Pathways via Promoter Binding to Mediate RBP4 Function

In Huh7 cells, Cut&Tag analysis was performed for BACH1, which directly binds to RBP4. The DNA targets bound by BACH1 were predominantly enriched in transcription start site (TSS) regions (Figure 2A). The genomic distribution of peaks across functional regions revealed that over 50% of the BACH1-bound genes were located within promoter regions, suggesting that most targets were directly regulated at promoter sites (Figure 2B). Analysis of peak distances relative to the TSS demonstrated that the majority of binding sites were located within 1 kb upstream and downstream regions of TSS, indicating that BACH1 binding is primarily concentrated near transcription initiation regions (Figure 2C).

**Fig 2.**
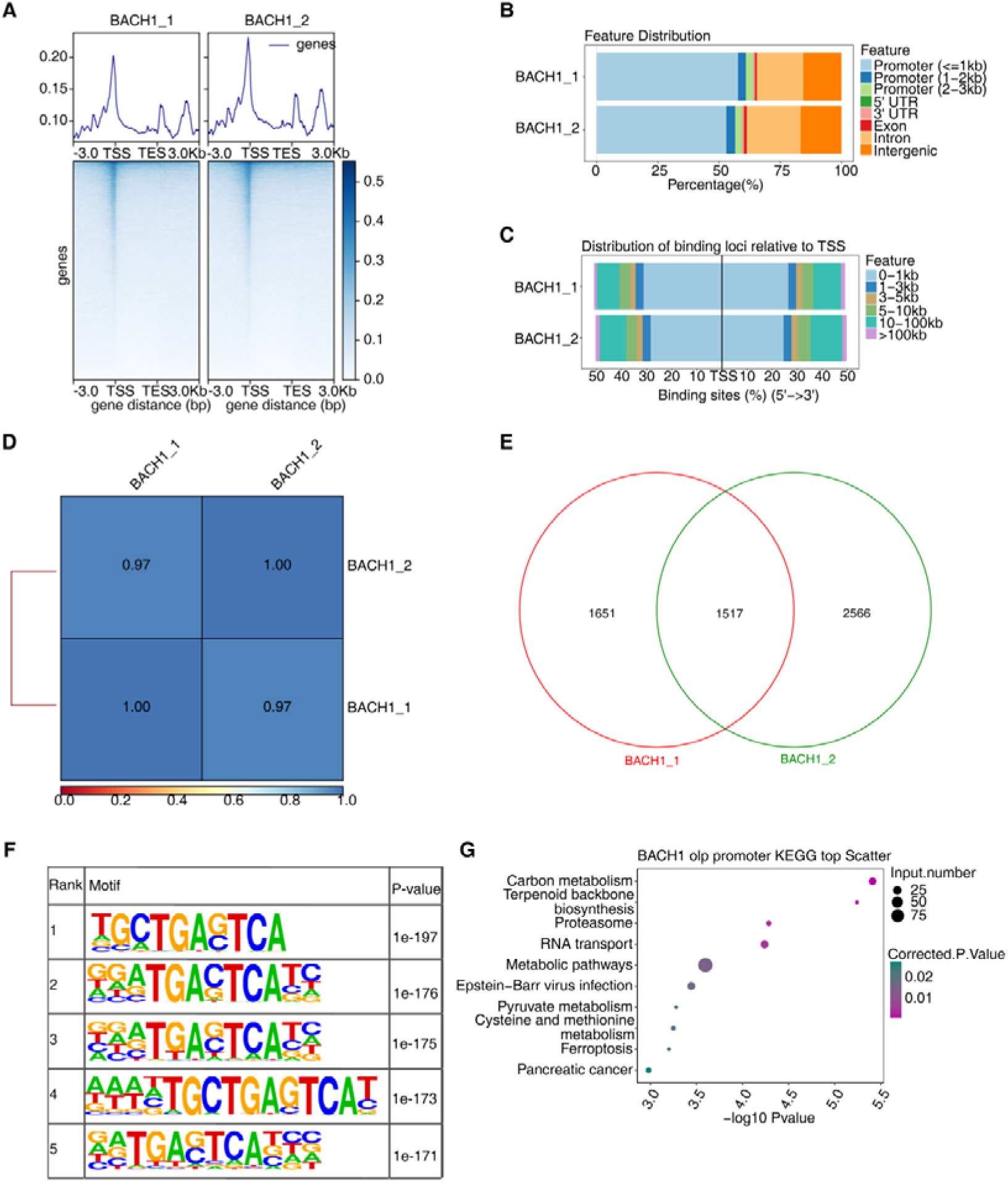
CUT&Tag analysis of BACH1 selectively binds DNAs in Huh7 cells. (A) Heatmaps of CUT&Tag signals around genomic binding regions within 3kb of gene. (B) Bars show the distribution of all samples in the functional regions of the peak genes. (C) Bars show the distance distribution of peak across all samples upstream and downstream of the TSS. (D) The correlation heatmap indicates the similarity of expression patterns between samples, and the closer the correlation coefficient is to 1, the higher the similarity of expression patterns between samples. (E) The Venn diagram merges Peak within groups (combined Peak lengths are 401 by default) and overlap analysis. (F) Top five motifs in the BACH1 genomic binding regions identified by the BACH1 CUT&Tag assay in Huh7 cells. (G) The KEGG enrichment analysis of the genes containing overlap peaks detected.

As a transcription factor, BACH1 may regulate downstream gene expression following its interaction with RBP4. Correlation analysis between two independent Cut&Tag experiments showed high reproducibility, with a correlation coefficient of 0.97 (Figure 2D). Overlap analysis of binding peaks between replicates identified 1,517 shared peaks (Figure 2E). Motif analysis revealed that both experiments identified the consensus binding motif TGACTCA (Figure 2F), consistent with previously reported BACH1 binding sequences(Warnatz *et al*, 2011). KEGG enrichment analysis of genes associated with the overlapping peaks demonstrated that the top enriched pathways included carbon metabolism, terpenoid backbone biosynthesis, pyruvate metabolism, cysteine and methionine metabolism, cellular maintenance and disease pathways, proteasome function, RNA transport, metabolic pathways, Epstein-Barr virus infection, ferroptosis, and pancreatic cancer. Among these, metabolic pathways showed the most significant enrichment (Figure 2G). Collectively, these results indicate that BACH1 binds to a large number of genes involved in metabolic processes in Huh7 cells, thereby playing a critical role in regulating cellular metabolism.

### 2.3 RBP4 Involved in the Transcription and Expression of Downstream Insulin Signaling Pathway-Related Genes by Binding to BACH1

To further investigate the potential target genes co-regulated by RBP4 and BACH1, a three-way overlap analysis was performed, integrating the differentially expressed genes (DEGs) from RNA-seq following RBP4 overexpression, the 1,090 BACH1-bound genes identified through Cut&Tag, and the predicted BACH1 target genes from three transcription factor databases. This analysis identified 397 genes potentially co-regulated by RBP4 and BACH1, including 334 previously known BACH1 targets and 63 newly identified targets in this study (Figure 3A). GO enrichment analysis of these 397 genes revealed significant enrichment in pathways related to DNA repair and the ubiquitin-dependent proteasomal protein catabolic process (Figure 3B). Considering the established roles of RBP4 and BACH1 in insulin resistance, glucose homeostasis, and lipid metabolism(Gliniak *et al*., 2017; Jin *et al*, 2023), we focused on pathways relevant to these biological processes. The analysis revealed significant enrichment of genes associated with the insulin receptor signaling pathway and carbohydrate metabolism. Among these, insulin receptor signaling pathway-related genes included GSK3B and PIK3R3, with detailed binding and expression profiles presented in Figure S1.

**Fig 3.**
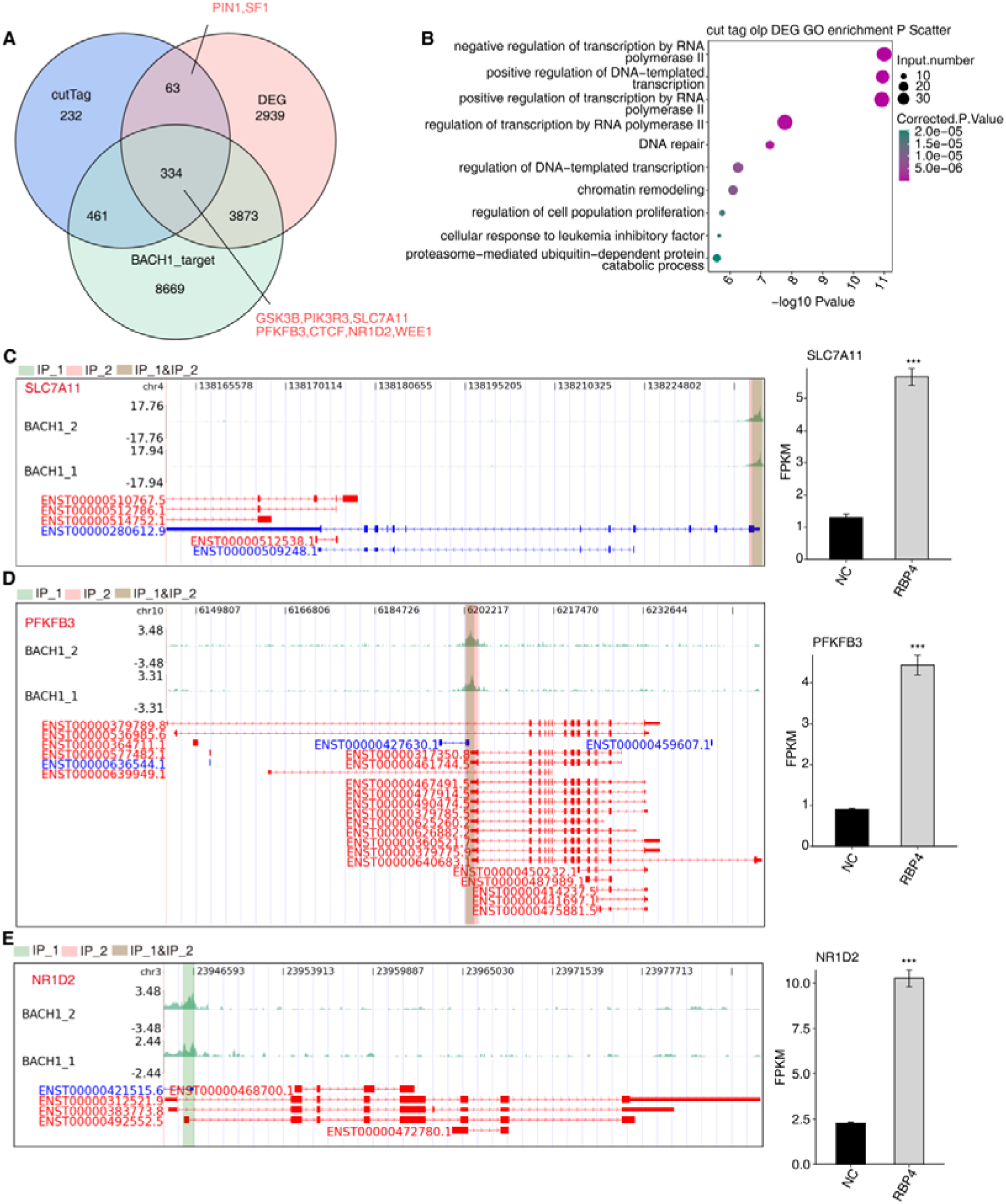
RBP4 regulated gene expression through directly binding its promoter by interact with BACH1. (A) Venn plots showing the overlap analysis of promoter peak genes with predicted BACH1 targets and differential genes. (B) Promoter peak functional enrichment analysis of overlap genes. (C-E) Left panel: CUT&Tag assay proved that BACH1 directly bound to SLC7A11, PFKFB3, and NR1D2 promoter region. IGV-sashimi plot showing the peaks reads and binding sites across DNA, the green panels represent the position of peaks. Reads distribution of bound gene is plotted in the up panel and the transcripts of each gene are shown below. Right panel: The schematic diagrams depict the structures of DEG. RNA-seq validation of SLC7A11, PFKFB3, and NR1D2 are shown at the bottom of the right panel. Error bars represent mean ± SEM.*** P-value < 0.001.

The 397 genes were ranked by their normalized log -transformed read count scores, and key targets were selected for further investigation. Known targets included SLC7A11, PFKFB3, and NR1D2 (Figure 3C–E), as well as CTCF and WEE1 (Figure S1), while PIN1 and SF1 were identified as novel targets in this study (Figure S2). Taken together, these findings demonstrate that RBP4 regulates the transcription and expression of downstream target genes, such as GSK3B and PIK3R3, by binding to BACH1, thereby participating in the regulation of insulin signaling and metabolic processes.

## Discussion

Recent studies have demonstrated that RBP4 not only functions as a crucial protein in the regulation of retinol metabolism but also serves as a key component in the modulation of hepatic circadian rhythm and metabolic homeostasis. Through systematic experimental analyses, the present study revealed that RBP4 directly interacts with the TF BACH1. This interaction contributes to the regulation of the insulin receptor signaling pathway and carbohydrate metabolism. These findings bridge an important knowledge gap regarding the molecular mechanisms through which RBP4 regulates hepatic metabolism.

Using Co-IP coupled with MS, 543 RBP4-interacting proteins were identified for the first time in Huh7 cells, including 19 TFs. Among these, BACH1 was selected as a key target for further investigation due to its established involvement in melatonin rhythm regulation(Li *et al*, 2022) and its known regulatory role in hepatic insulin signaling(Jin *et al*., 2023). Previous reports have indicated that BACH1 can participate in protein–protein interactions through its BTB domain(Jin *et al*., 2023), and our mass spectrometry results confirmed a direct interaction between RBP4 and BACH1. However, whether this binding is mediated via the BTB domain requires further experimental verification. RBP4, as an adipokine, binds to STRA6(Gliniak *et al*., 2017) and induces internal phosphorylation of the STRA6 receptor, thereby triggering the JAK2/STAT5 signaling cascade and promoting SOCS3 expression(Bako *et al*, 2019; Shi *et al*, 2006; Steinhoff *et al*, 2021). SOCS3 subsequently reduces IRS1 protein levels, leading to decreased insulin-stimulated glucose uptake. The RBP4-BACH1 axis identified in the present study may represent an independent, complementary, or cross-talking mechanism distinct from this canonical STRA6-mediated pathway. This finding adds transcriptional regulation as a new dimension to the multifaceted biological functions of RBP4. Although RBP4 is a secreted protein, hepatocytes serve as the primary source of circulating RBP4. It is plausible that, similar to other specialized secretory proteins such as fibroblast growth factor-2(Nickle *et al*, 2024) and interleukin-1α(Rider *et al*, 2013), RBP4 may also exert intracellular functions. In addition, RBP4 might undergo nuclear translocation or bind to immature BACH1 precursors in the endoplasmic reticulum/Golgi apparatus. This intriguing possibility warrants further investigation. Future studies should explore whether specific RBP4 isoforms, post-translational modifications (e.g., phosphorylation), or binding partners determine its fate—whether it is directed toward the secretory pathway or retained intracellularly to interact with factors such as BACH1.

The identification of BACH1 as a core TF specifically bound by RBP4 provides new insights into the nuclear functions of RBP4, which has traditionally been recognized primarily as a retinol transport protein. This finding suggests that RBP4 may indirectly regulate metabolic processes through its interactions with transcription factors.

CUT&Tag analysis demonstrated that BACH1 binding sites were predominantly distributed within promoter regions (0–1 kb, >50%), consistent with its role as a TF capable of mediating downstream transcriptional regulation after binding to RBP4. Furthermore, the identified binding motif of BACH1, TGACTCA, closely resembled the recognition site of CREB(Sassone-Corsi *et al*, 1990). Given that the CREB/CRTC2 transcriptional complex has been shown to regulate hepatic circadian rhythms(Sun *et al*, 2015) and glucose homeostasis(Morgan *et al*, 2024), these findings suggest that RBP4 may engage competitively or cooperatively with the CREB/CRTC2 pathway through its interaction with BACH1. In addition, CUT&Tag analysis identified core clock genes, such as NR1D2, among the BACH1 target genes, which were significantly enriched in metabolic pathways including carbon metabolism, pyruvate metabolism, and insulin signaling. Notably, previous studies have demonstrated that RBP4 expression exhibits circadian rhythmicity(Ma *et al*, 2016), and disruption of circadian rhythms leads to marked upregulation of RBP4 expression(Wilms *et al*, 2019). Taken together, these findings suggest that RBP4 may function as a pivotal node linking hepatic circadian rhythm regulation with glucose homeostasis.

Integrated analysis of RNA-seq, CUT&Tag, and three TF databases identified 397 genes co-regulated by RBP4 and BACH1. Among these, molecules such as GSK3B and PIK3R3 exhibited dual regulatory roles in circadian rhythm modulation and insulin signaling. Specifically, GSK3B not only regulates core clock components(Yuan *et al*, 2021) but also modulates glucose and lipid metabolism(Yan *et al*, 2023). Pik3r3-knockdown mice have been shown to exhibit impaired glucose tolerance(Zhu, 2023), and PIK3R3 influences clock gene expression through the PI3K/AKT signaling pathway(Khezri *et al*, 2024). In conjunction with the downstream targets of the RBP4–BACH1 interaction identified in this study—including known targets such as PFKFB3 and NR1D2, as well as newly identified targets such as PIN1—these results suggest that the RBP4–BACH1 axis may function through multiple mechanisms: direct regulation of clock-related genes such as NR1D2, modulation of insulin signaling components such as PFKFB3, and regulation of protein homeostasis via molecules such as PIN1. Collectively, these processes form a multilayered regulatory network governing hepatic metabolism.

This study demonstrates that RBP4-BACH1 interaction participates in hepatic metabolic reprogramming by influencing BACH1-mediated transcriptional regulation, thereby providing new mechanistic insights into how the RBP4–BACH1 axis integrates circadian rhythms with glucose metabolic homeostasis. Future studies should focus on generating liver-specific RBP4–BACH1 double-knockout animal models to validate the in vivo functions of this axis. In addition, the development of small-molecule modulators targeting their interaction interface should be prioritized. Furthermore, the clinical relevance of RBP4 and BACH1 expression profiles in glucose metabolism disorders among shift workers warrants detailed investigation.

There are some limitations of this study. First, the use of Huh7 cells with RBP4 overexpression, while providing valuable mechanistic insights, may not fully recapitulate physiological conditions in vivo. Overexpression systems can potentially force protein-protein interactions that occur only at supraphysiological concentrations or promote non-specific binding, leading to artifacts that may not reflect endogenous regulatory mechanisms. Second, as a transformed cell line, Huh7 cells may exhibit altered metabolic and transcriptional programs compared to primary human hepatocytes, which could influence the generalizability of our findings. Therefore, the results require further validation in various physiological or disease-related hepatocyte models such as primary hepatocytes and liver organoids, as well as animal models, to confirm the role of the RBP4-BACH1 axis in metabolic regulation in vivo. Third, although our multi-omics integration supports a functional interaction between RBP4 and BACH1, the precise subcellular localization of this interaction—whether cytoplasmic, nuclear, or both—remains to be determined. It is noteworthy that RBP4 has traditionally been characterized as a secreted adipokine involved in retinol transport; however emerging evidence supports the concept of moonlighting proteins which perform multiple context-dependent functions beyond their canonical roles. In this paradigm, RBP4 may exert intracellular functions through interaction with transcription factors such as BACH1, particularly under conditions of metabolic stress or circadian disruption.

To further elucidate this mechanism, future studies should elucidate the RBP4-BACH1 axis under physiologically relevant conditions, future studies should focus on detailed mechanistic and transport analyses. First, the response to retinol stimulation—including the effects of vitamin A supplementation on RBP4 secretion, subcellular localization, and its interaction with BACH1—requires systematic evaluation. Subcellular fractionation coupled with western blotting should be employed to analyze the distribution of both endogenous and overexpressed RBP4 in the cytoplasmic, nuclear, and endoplasmic reticulum fractions. Second, immunofluorescence confocal microscopy should be used to observe the co-localization of endogenous RBP4 and BACH1 under various conditions, including normal culture, ER stress, and retinol stimulation. This will help determine whether a nuclear pool of RBP4 can be detected under physiological conditions and clarify the processing pathway and efficiency of RBP4 in hepatocyte models. Third, CRISPR/Cas9-mediated endogenous tagging can be utilized to investigate the localization and interactions of RBP4 at endogenous expression levels, thereby avoiding artifacts introduced by transient overexpression. To enhance translational significance, validation of RBP4 expression profiles in clinical samples, such as those from shift workers, is also necessary. Finally, the specific molecular mechanisms by which BACH1 regulates clock-related genes such as GSK3B warrant further experimental investigation.

This study identified BACH1 as a key transcription factor specifically interacting with RBP4. RBP4/BACH1 co-regulated 63 metabolism-related genes, including SF1 and PIN1, which were newly identified as targets associated with circadian rhythm. RBP4 modulates insulin signaling pathways through BACH1, thereby driving glucose metabolic reprogramming and offering novel therapeutic targets for the prevention and treatment of circadian rhythm disruption-associated glucose metabolism disorders.

## Methods

### Structured Methods - Reagents and Tools Table

| Reagent/Resource | Reference or Source | Identifier or Catalog Number |
| --- | --- | --- |
| **Experimental Models** |  |  |
| Huh7 cell line (human hepatoma) | Procell Life Science & Technology Co., Ltd. | N/A |
| **Recombinant DNA** |  |  |
| pcDNA3.1-3×Flag-RBP4 | Youbio Biotech (Changsha, PRC) | NM_006744 |
| siRNA targeting RBP4 | Youbio Biotech | This study |
| **Antibodies** |  |  |
| Anti-RBP4 | Proteintech | 66104-1-Ig |
| Anti-RBP4 | Proteintech | 11774-1-AP |
| Anti-BACH1 | Proteintech | 14018-1-AP |
| Anti-GAPDH | AtaGenix | ATPA00013Rb |
| Anti-H3 | Proteintech | 17168-1-AP |
| HRP-conjugated goat anti-rabbit IgG | Proteintech | SA00001-2 |
| HRP-conjugated goat anti-mouse IgG | Proteintech | SA00001-1 |
| Normal rabbit IgG | ABclonal | AC005 |
| **Oligonucleotides and other sequence-based reagents** |  |  |
| qPCR primers for human GAPDH | This study | N/A |
| qPCR primers for human RBP4 | This study | N/A |
| **Chemicals, Enzymes and other reagents** |  |  |
| DMEM | Gibco / ThermoFisher | N/A |
| Fetal bovine serum (FBS) | Gibco / ThermoFisher | N/A |
| Penicillin/streptomycin | Gibco / ThermoFisher | N/A |
| Lipofectamine 2000 | Invitrogen / ThermoFisher | N/A |
| TRIzol reagent | Invitrogen / ThermoFisher | 15596026 |
| Hifair® II 1st Strand cDNA Synthesis SuperMix | Yeasen Biotec | 11123ES60 |
| Hieff® qPCR SYBR Green Master Mix | Yeasen Biotec | 11202ES08 |
| RIPA lysis buffer | Beyotime | N/A |
| BCA protein assay kit | BeyotimeBeyotime | N/A |
| PVDF membrane | N/A | N/A |
| ECL detection kit | peptbio | N/A |
| NP-40 lysis buffer | This study | N/A |
| Protease inhibitor cocktail | Bimake China | B14001 |
| Protein A/G magnetic beads | Thermo Scientific | 26162 |
| Silver staining kit | Beyotime | P0017S |
| CUT&Tag assay kit | Yeasen | 12598ES12 |
| DNase I | N/A | N/A |
| Qubit™ RNA Broad Range Assay Kit | Life Technologies | Q10210 |
| KCTM Stranded mRNA Library Prep Kit | Wuhan Seqhealth Co., Ltd. | DR08402 |
| **Software** |  |  |
| MaxQuant (1.6.6.0) | Max Planck Institute | https://www.maxquant.org |
| Andromeda search engine | Integrated in MaxQuant | N/A |
| Bowtie2 (v2.4.4) | Johns Hopkins University | http://bowtie-bio.sourceforge.net |
| Sambamba (0.8.1) | Open source | https://lomereiter.github.io/sambamba |
| Samtools | Open source | http://www.htslib.org |
| MACS2 (v2.2.7.1) | Open source | https://github.com/macs3-project/MACS |
| HOMER (motif analysis) | UCSD | http://homer.ucsd.edu |
| KOBAS 2.0 server (functional enrichment) | Peking University | http://kobas.cbi.pku.edu.cn |
| GraphPad Prism (version 8.0) | GraphPad Software | https://www.graphpad.com |
| **Other** |  |  |
| NanoDrop™ OneC spectrophotometer | Thermo Fisher Scientific | N/A |
| Qubit 3.0 fluorometer | Life Technologies | N/A |
| NovaSeq 6000 / NovaSeq X Plus | Illumina | N/A |
| UltiMate 3000 RSLCnano system coupled with Q Exactive HF MS | Thermo Fisher Scientific | N/A |
| Nanospray Flex ion source | Thermo Fisher Scientific | N/A |

### 1.1 Cell Culture

Huh7 cell lines were obtained from Procell Life Science & Technology Co., Ltd. (China) and cultured in Dulbecco’s Modified Eagle Medium (DMEM) supplemented with 10% fetal bovine serum (FBS), 100 μg/mL streptomycin, and 100 U/mL penicillin at 37°C in a humidified atmosphere containing 5% CO□.

### 1.2 Plasmid and Transfections

The pcDNA3.1-3×Flag-RBP4 plasmid was purchased from Youbio Biotech (NM_006744, Changsha PRC). Plasmid transfection into Huh7 cells was performed using Lipofectamine 2000 (Invitrogen, Carlsbad, CA, USA) in accordance with the manufacturer’s protocol for RBP4 overexpression. Transient RBP4 knockdown in Huh7 cells was constructed by transfection with specifically synthesized siRNA by Youbio Biotech (sense 5’-AUGGCAGAUCAGAAAGAAAUU-3’, antisense 5’-UUUCUUUCUGAUCUGCCAUUU-3’). Transfected cells were harvested 48 hours post-transfection for RT-qPCR, Western blot analyses or RNA-seq.

### 1.3 RT-qPCR

Total RNA was extracted from cells using TRIzol reagent following the manufacturer’s instructions (15596026, Invitrogen). Complementary DNA (cDNA) synthesis was performed using the Hifair® II 1st Strand cDNA Synthesis SuperMix (11123ES60, Yeasen Biotec). Quantitative PCR was carried out using Hieff® qPCR SYBR Green Master Mix (11202ES08, Yeasen Biotec) to quantify the expression levels of target genes. Primer sequence (5’ −3’) Homo GAPDH F: TCAAGAAGTGGTGAAGCGG R: TCAAAGGTGGAGGTGGGT. Homo RBP4 F: GATGACCACTGGATCGTCGA R: TGCCTACAATCTTTGCGC.

### 1.4 Western Blotting and Antibodies

Cells were harvested, and total proteins were extracted using RIPA lysis buffer. Following centrifugation, the supernatant was collected, and the protein concentration was determined using a BCA assay kit (Beyotime). The extracted proteins were separated by SDS–PAGE and transferred onto a PVDF membrane. The membrane was blocked with 5% skim milk and subsequently incubated with the following primary antibodies: anti-RBP4 (1:2000, 66104-1-Ig, proteintech), anti-BACH1 (1:2000, 14018-1-AP, proteintech), anti-GAPDH (1 : 5000, ATPA00013Rb, AtaGenix) and anti-H3 (1:2000, 17168-1-AP, proteintech). After washing, the membrane was incubated with horseradish peroxidase (HRP)-conjugated secondary antibodies, either HRP-conjugated goat anti-rabbit IgG (1:5000, SA00001-2, Proteintech) or HRP-conjugated goat anti-mouse IgG (1 : 5000, SA00001-1, Proteintech). The PVDF membranes were cut before hybridization with antibodies. The original images that can be used for full-length blotting were provided in Supplementary Figure S3. Protein bands were visualized using an enhanced chemiluminescence (ECL) detection kit (peptbio), and signal intensities were quantified.

### 1.5 RNA Extraction and Sequencing

Total RNA was extracted from Huh7 cells using TRIzol reagent (Invitrogen, cat. NO 15596026). Genomic DNA contamination was removed by DNase I digestion following RNA extraction. RNA purity was assessed by measuring the A260/A280 ratio using a NanoDrop™ OneC spectrophotometer (Thermo Fisher Scientific Inc), and RNA integrity was verified by 1.5% agarose gel electrophoresis. Qualified RNA samples were quantified using a Qubit 3.0 fluorometer with the Qubit™ RNA Broad Range Assay Kit (Life Technologies, Q10210).

A total of 2 μg of RNA was used for stranded RNA-seq library preparation using the KCTM Stranded mRNA Library Prep Kit for Illumina (Catalog NO. DR08402, Wuhan Seqhealth Co., Ltd. China), following the manufacturer’s instructions. PCR products ranging from 200 to 500 bp were enriched, quantified, and sequenced on the NovaSeq 6000 platform (Illumina) using the PE150 mode.

### 1.6 Co-IP and MS

Co-IP assays were conducted by Wuhan Ruixing Biotechnology Co., Ltd (http://www.rxbio.cc). Cells were lysed on ice for 30 minutes in NP-40 buffer (50 mM Tris (pH 7.4), 150 mM NaCl, 20 mM EDTA and 1% NP-40) supplemented with a freshly added protease inhibitor cocktail (B14001, Bimake China). After centrifugation at 13,200 rpm for 10 minutes at 4°C, the supernatant was collected and incubated overnight at 4°C with rotation, with either 10 μg of anti-RBP4 antibody (immunoprecipitation group, 11774-1-AP, Proteintech) or IgG (negative control, AC005, ABclonal, China). Subsequently, 50 μL of protein A/G magnetic beads (26162, Thermo Scientific, USA) was added, and the mixture was rotated for 2 hours at 4°C.

The beads were washed three times with wash buffer (50 mM Tris, pH 7.4; 150 mM NaCl; 1 mM EDTA; supplemented with protease inhibitors). Bound proteins were eluted and subjected to SDS–PAGE. The gel was processed according to the manufacturer’s instructions for silver staining (P0017S, Beyotime, China), and all protein products were analyzed by MS (UltiMate 3000 RSLCnano system coupled on-line with Q Exactive HF MS through a Nanospray Flex ion source, Thermo).

### 1.7 MS Data Analysis

MS raw data were processed using MaxQuant software (1.6.6.0) with the Andromeda search algorithm.

The spectral data were searched against the human protein sequence database downloaded from UniProt (20230619), with the following parameters: variable modifications were set as Carbamidomethyl (C)-57.021464, Oxidation (M)-15.994915, Acetyl (Protein N-term)-42.010565, and Deamidation (NQ)-0.984016. Carbamidomethyl (C)-57.021464 was also included as a fixed modification. Trypsin/P was specified as the digestion enzyme, allowing a maximum of two missed cleavages. The peptide mass tolerances for the first and main searches were set to 20 ppm and 4.5 ppm, respectively, and the fragment match tolerance was set to 20 ppm.

Proteins that could not be distinguished based on unique peptides were merged into a single protein group by MaxQuant. The search results were filtered at a 1% false discovery rate (FDR) at both the peptide and protein levels. Proteins identified as reverse sequences, potential contaminants, or identified by site only were excluded. The “proteinGroups.txt” file was used for protein-level analysis. Label-free quantification (LFQ) intensities were log -transformed, and missing values were imputed using values from a normal distribution. Proteins significantly enriched in anti-RBP4 antibody samples compared with isotype controls were selected based on log fold-change thresholds (two-tail t-test p-value < 0.05).

### 1.8 CUT&Tag Sequencing and Data Processing

CUT&Tag assays were performed in Huh7 cells to profile the chromatin binding of BACH1. A total of 1×10 cells were collected per assay, and 1 μL of anti-BACH1 antibody (14018-1-AP, proteintech) was used according to the manufacturer’s instructions (Yeasen, Cat. No. 12598ES12). DNA libraries were prepared by PCR amplification as specified in the protocol (Yeasen). Library fragments ranging from 200 to 500 bp were enriched, quantified, and sequenced on an Illumina NovaSeq X Plus system using the PE150 mode.

Paired-end reads were aligned to the GRCh38 human genome using Bowtie2 (v2.4.4) with the parameters: --local, --very-sensitive, --no-mixed, --no-discordant, -I 10, -X 700. Duplicate reads were removed using Sambamba (0.8.1). Reads mapped to canonical chromosomes were retained, and both uniquely and multi-mapped reads were extracted using Samtools. The BAM files were sorted and indexed with Samtools, and peak calling was performed using MACS2 (v2.2.7.1). Narrow peaks were called with the parameters -f BAMPE --keep-dup all, and broad peaks were called using --broad --broad-cutoff 0.1 -f BAMPE --keep-dup all. Processed BAM files were used to extract read counts within identified peaks. Enriched binding motifs within peaks were analyzed using HOMER (Hypergeometric Optimization of Motif EnRichment) software(Heinz *et al*, 2010).

### 1.9 TFs

A catalogue of 1796 transcription factors of human was retrieved and combined from two previous reports(Hu *et al*, 2019; Lambert *et al*, 2018).

### 1.10 Functional Enrichment Analysis

To identify functional categories of peak-associated genes (target genes), Gene Ontology (GO) terms and Kyoto Encyclopedia of Genes and Genomes (KEGG) pathways were analyzed using the KOBAS 2.0 server(Xie *et al*, 2011). The hypergeometric test combined with the Benjamini–Hochberg FDR correction was applied to determine statistically significant enrichment.

### 1.11 Statistical Analyses

Data obtained from at least three independent experiments are presented as mean ± standard deviation (SD). Statistical analyses were performed using Student’s t-test for two-group comparisons or two-way analysis of variance (ANOVA) for comparisons among multiple groups. A p-value of < 0.05 was considered statistically significant (*p < 0.05; **p < 0.01; ***p < 0.001). All graphical representations were generated using GraphPad Prism software (version 8.0, San Diego, CA, USA).

## Data availability

The source data of this paper are available in the following databases:

RNA-Seq data: Genome Sequence Archive in National Genomics Data Center <u>HRA016389(</u>https://ngdc.cncb.ac.cn/gsa-human)

Co-IP followed by MS data: Genome Sequence Archive in National Genomics Data Center <u>HRA016411</u> (https://ngdc.cncb.ac.cn/gsa-human)

## Authors contributions

Li Wang: Writing – original draft, Supervision, Project administration. Qi Ma: Writing – original draft. ChenYao: Writing – original draft. Chunmei Wu: Visualization. Baoping Guo: Visualization. Mayire Nuermaimaiti: Investigation. Yinxia Su: Data curation. Binbin Fang: Validation. Yi Zheng: Investigation. Xiangrui Chen: Investigation. Lijuan He: Writing - review & editing. Guoli Du: Writing - review & editing.

## Acknowledgements

This project was supported by the State Key Laboratory of Pathogenesis, Prevention and Treatment of High Incidence Diseases in Central Asia Fund (SKL-HIDCA-2024-13), the Youth Science and Technology Elite Talent Program, Xinjiang Medical University (XYD2024Q07), Tianshan Elite — High-Level Medical Talents Training Program of the Health Commission of Xinjiang Uygur Autonomous Region (TSYC20230B079).

We acknowledge the technicians from Wuhan Ruixing Biotechnology Co. for their help in generating data presented in this study.

---

**Fig S1.**
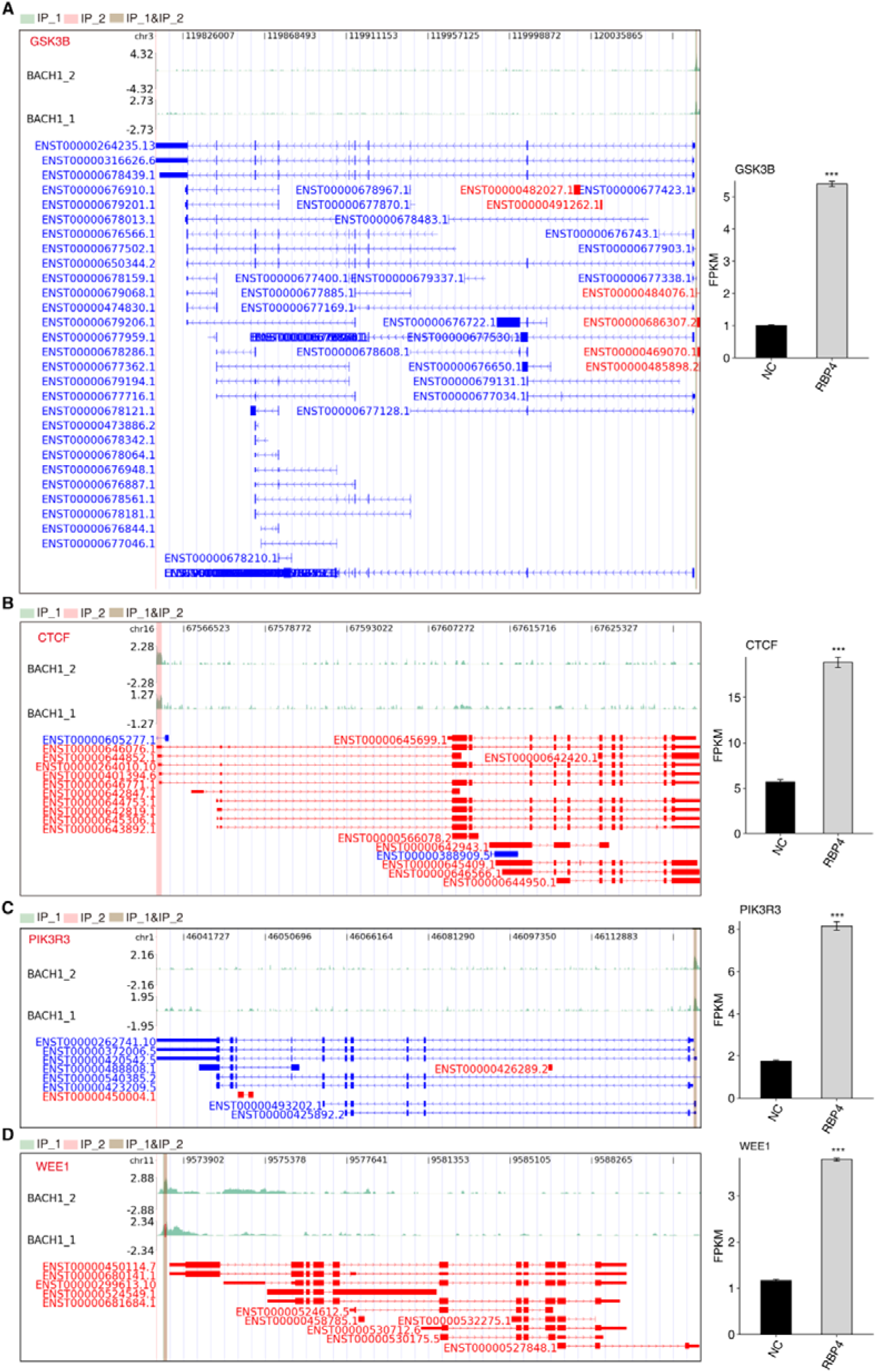
(A-D) Left panel: CUT&Tag assay proved that BACH1 directly bound to GSK3B, CTCF, PIK3R, and WEE1 promoter region. IGV-sashimi plot showing the peaks reads and binding sites across DNA, the green panels represent the position of peaks. Reads distribution of bound gene is plotted in the up panel and the transcripts of each gene are shown below. Right panel: The schematic diagrams depict the structures of DEG. RNA-seq validation of GSK3B, CTCF, PIK3R, and WEE1 are shown at the bottom of the right panel. Error bars represent mean ± SEM. *** P-value < 0.001.

**Fig S2.**
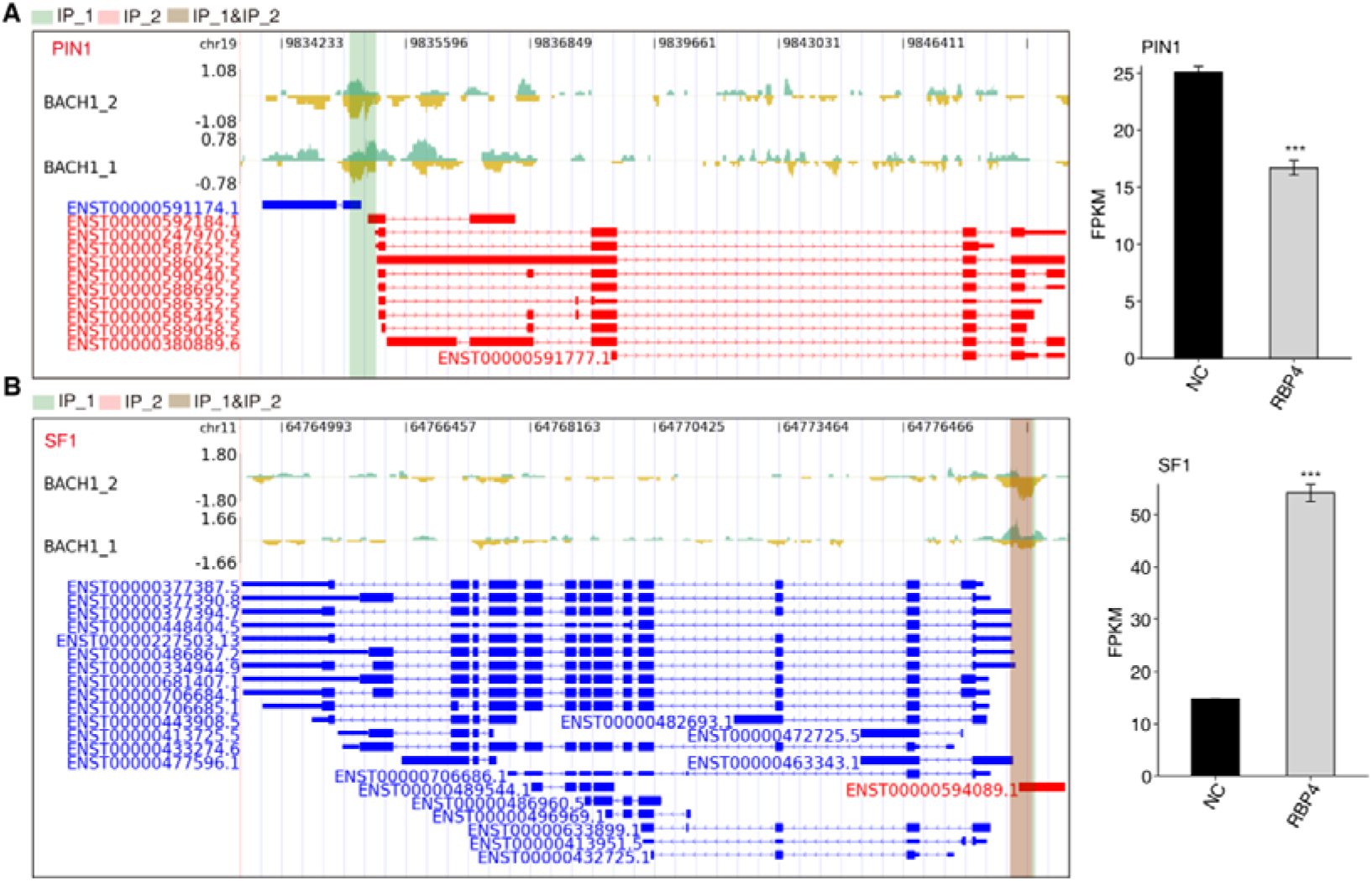
(A-B) Left panel: CUT&Tag assay proved that BACH1 directly bound to PIN1 and SF1 promoter region. IGV-sashimi plot showing the peaks reads and binding sites across DNA, the green panels represent the position of peaks. Reads distribution of bound gene is plotted in the up panel and the transcripts of each gene are shown below. Right panel: The schematic diagrams depict the structures of DEG. RNA-seq validation of PIN1 and SF1 are shown at the bottom of the right panel. Error bars represent mean ± SEM.*** P-value < 0.001.

**Fig S3.**
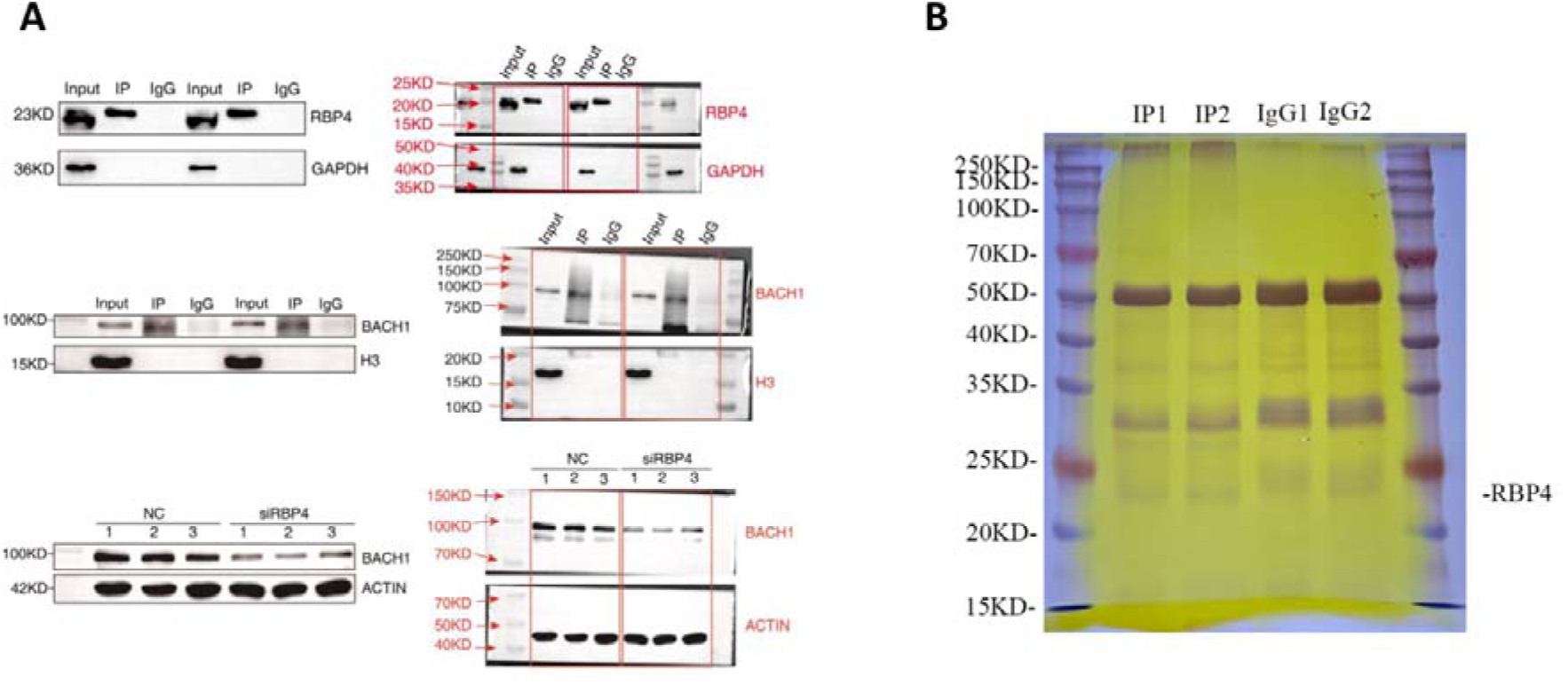
(A) The original images of blots exhibited in Figure 1H and Figure 1I, as well as the replicates. The blots were cut at 35 kDa prior to hybridization with antibodies. (B) Silver staining detection of RBP4-interacting proteins.

